# Liberality is More Explainable than PCA of Transcriptome for Vertebrate Embryo Development

**DOI:** 10.1101/2024.09.19.613970

**Authors:** Norichika Ogata

## Abstract

Dimensional reduction approaches of genome scale data are widely used. Liberality is one of the approaches which is a quantitative index of cellular differentiation and dedifferentiation. Here, we analyzed a trend of liberality of time course transcriptome data on vertebrate embryo development. Historically annotated embryo developmental stages matched changes in the trend of liberality. Analyzing liberality of various biological phenomena would be beneficial.

## I. Introduction

Genome scale raw data does not fit to recognise and understand or find something for us. Most studies analyze and summarize these data using dimensional reduction methods. In this decade, the most major method used for genome expression data would be the primary component analysis (PCA). A disadvantage of the PCA is the meaninglessness of values in the PCA such as PC1, PC2. In the PCA, we place the degree of similarity of the differences in the samples used in the analysis among the samples used in the analysis. In ecology, the PCA is used to estimate the beta diversity of samples[1]. Along with the beta diversity, there is the concept of the alpha diversity; beta diversity is the quantification of diversity inter samples, and alpha diversity is the quantification of diversity intra the sample. Therefore, we need several samples to estimate beta diversity and we can estimate alpha diversity from only one sample. To estimate the alpha diversity of genome expression data, Shannon’s information entropy of numerically converted RNA-seq data (with reference genome/transcriptome sequences, alignment/mapping processes, and tag-counting/assembling processes) [2] and Lempel-Ziv complexity directly measured from sequence data of RNA-seq [3] have being used. In a previous study, the biological meaning of alpha diversity of genome expression data was discovered; the cellular differentiation makes the value low and the cellular dedifferentiation makes the value high [4]. After that several studies have reached the same conclusion [5–8]. To represent a degree of cellular differentiation/dedifferentiation, we started using a term “liberality”. Liberality is the quantified value of the degree of cellular differentiation/dedifferentiation in the cellular biological context [9–13]. It can be easily distinguished from some diversity; there is much diversity such as interspecific diversity[14], intraspecific diversity, cellular heterogeneity[15], environmental diversity and more. Here, we will apply liberality to a genome expression data set of embryonic development. The data set was published in a study[16]. In the study, the PCA analysis was performed. We found no evidence to provide a reasonable explanation for the annotation of continuous phenomena cut into stages from the analysis. In this study, we compared numbers of inflection points in trends of liberality and values of PCA. We are going to explain the historically annotated embryo developmental stages with the genome scale expression data sets.

**Fig. 1.**
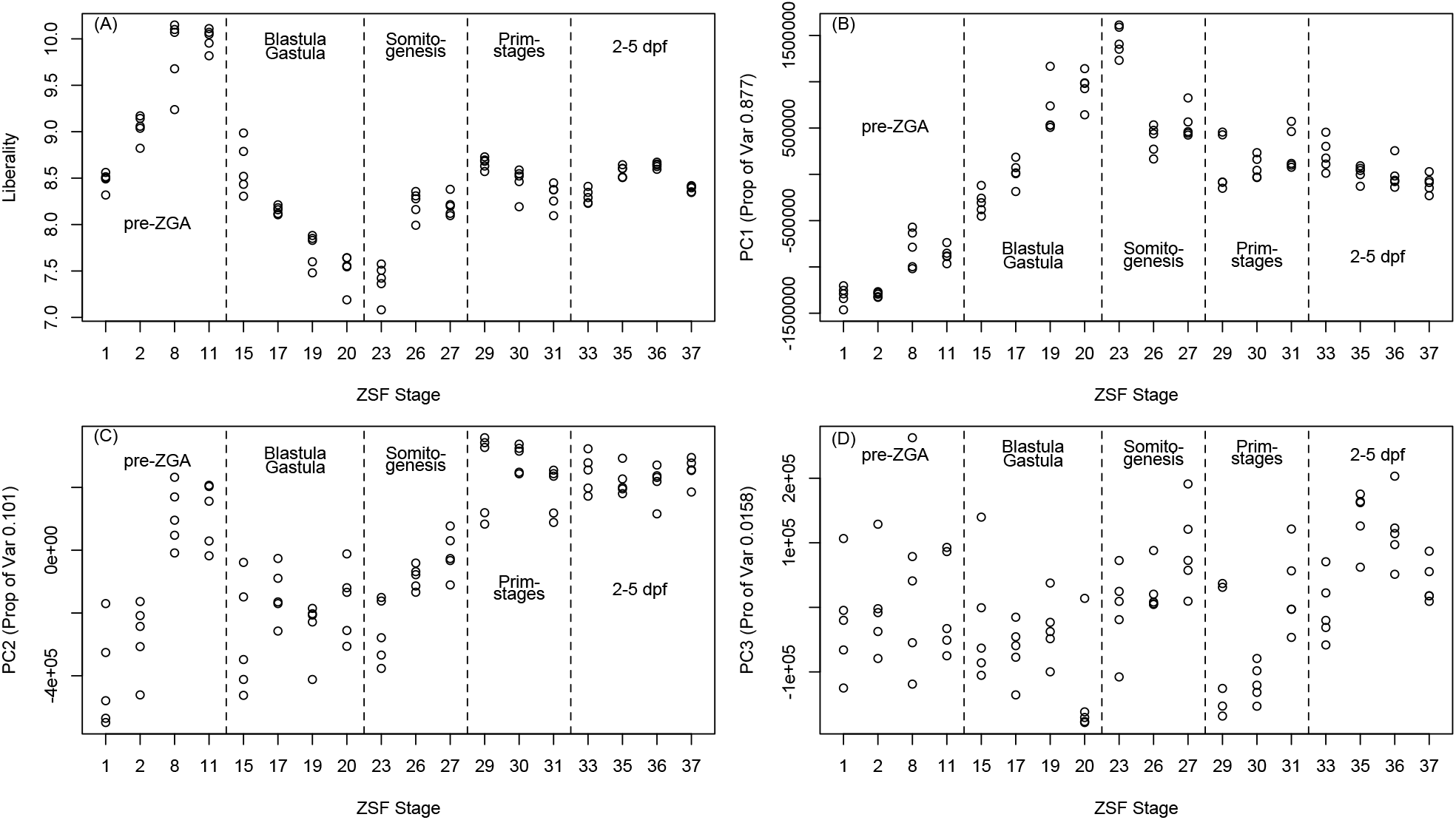
Liberality and Primary Components of mRNA expression time course of embryonic development in zebrafish. (A) Liverality vs ZSF stage. (B) Primary component 1 vs ZSF stage. (C) Primary component 2 vs ZSF stage. (D) Primary component 3 vs ZSF stage.

## II. Result s and Discussion

Liberality and primary components were estimated from the previously published mRNA expression time course of embryonic development in zebrafish, Danio rerio [16]. The samples which are the source of the data were from 0 hours post fertilization to 5 days post fertilization and were taken 18 times. Biological replication was five. In total, 90 RNA-seq were performed. We obtained the supplemental data 2 of the publication, RNA-seq count data in tsv format.

We plotted the liberality and primary components with the Zebrafish Stage Ontology (ZSF) stages [17]. In the plot of the liberality, we could find at least two inflection points in trends between ZSF0000011-ZSF0000015 and between ZSF0000023-ZSF0000026. In the plot of primary components (PC) 1, we could find only a single inflection point in trends between ZSF0000023-ZSF0000026. In the plots of PC2 and PC2, there is so much variability, we could not find any inflection points. Between ZSF0000011-ZSF0000015, embryos change their stage from the pre-GZA to the blastula/gastula. Between ZSF0000020-ZSF0000023, embryos change their stage from the blastula/gastula to the somitogenesis. Liberality was more explainable than PCA of transcriptome for the annotation of embryonic development..

## III. Materials and methods

The RNA-seq count data of mRNA expression time course of embryonic development in zebrafish were obtained from the publication. The PCA analysis was performed using prcomp command on R version 4.4.1. The liberality was estimated as previously reported [10]. The implementation of the calculation method for liberality using the R language [18] is as follows.

~~~
entroshannon<-function(x){
x2<-x/sum(x)
x3<-x2*log2(x2)
x4<-x3[!is.na(x3)]
-sum(x4)
}
~~~

